# Beating your Neighbor to the Berry Patch

**DOI:** 10.1101/2020.11.12.380311

**Authors:** Alan R. Rogers

## Abstract

Foragers often compete for resources that ripen (or otherwise improve) gradually. What strategy is optimal in this situation? It turns out that there is no optimal strategy. There is no evolutionarily stable strategy (ESS), and the only Nash equilibrium (NE) is unstable: strategies similar to the NE can always invade. But in spite of this instability, the NE is predictive. If harvesting attempts are costly or there are many competitors, the process tends to remain near the unstable NE. In this case, the resource often goes unharvested. Harvesting attempts—when they happen at all—usually occur when the resource is barely ripe enough to offset costs. The more foragers there are, the lower the chance that the resource will be harvested and the greater its mean value when harvested. This counterintuitive behavior is exhibited not only by theoretical models and computer simulations, but also by human subjects in an experimental game.

## 1 Introduction

Every summer, my backyard witnesses a conflict between humans and birds, all of whom wish to eat the same strawberries. Those who wait until the berries are ripe eat none, for by then the others have come and gone. All of us eat sour berries or none at all, and none of us are happy about it.

Such interactions must be common in nature. They occur whenever

1. Several individuals compete for the same resource.
2. The resource improves in value over time.
3. Some cost is involved in attempting to harvest the resource whether one succeeds in harvesting it or not.
4. Harvesting the resource ruins it for those who come later.

I know of three examples from ethnography. (1) The Barí are a horticultural people in Venezuela whose fishing methods have been described by Bennett [1]. The Barí often fish by dumping poison into pools, where the stream is deep and slow. This kills most or all of the fish. New fish enter the pool only slowly, so that the pool improves in value over a period of weeks. The Barí defend territories, but different villages nonetheless exploit the same pools. The Barí therefore face a dilemma. If they wait until the pool is full of fish, another village may exploit the pool first. But if they go too soon, the pool is hardly worth exploiting. (2) The Hadza are a foraging people in Tanzania [3]. Like most tropical foragers, they enjoy honey. Hives improve in quality during the spring and summer, so it is best not to exploit them too soon. But the Hadza also compete for honey not only with each other but also with birds and badgers. Humans and badgers both destroy hives when they exploit them, so little is left for subsequent foragers. (3) The Aché are a population in Paraguay whose economy involves both foraging and gardening. Kim Hill (personal communication) tells me that garden products are seldom allowed to ripen because children roam the gardens foraging for themselves. Parents who waited for the produce to ripen would harvest nothing. Hill has worked with the Aché for decades but has yet to eat a ripe watermelon.

These examples show that the interaction in my back yard is not an isolated example. It illustrates a problem that must have confronted our ancestors for a very long time. Thus, it makes sense to ask what strategy would have been favored by natural selection. Below, I introduce a model that answers this question. First, however, I motivate the theory by showing how real people respond to similar dilemmas in classroom experiments.

## 2 A classroom experiment

Subjects were recruited from undergraduate anthropology classes, and the experiment was approved by the Institutional Review Board of the University of Utah. Subjects interacted with each other via a computer program, which provided instructions, calculated scores, and kept track of each subject’s choices. The screen is shown in figure 1.

**Figure 1.**
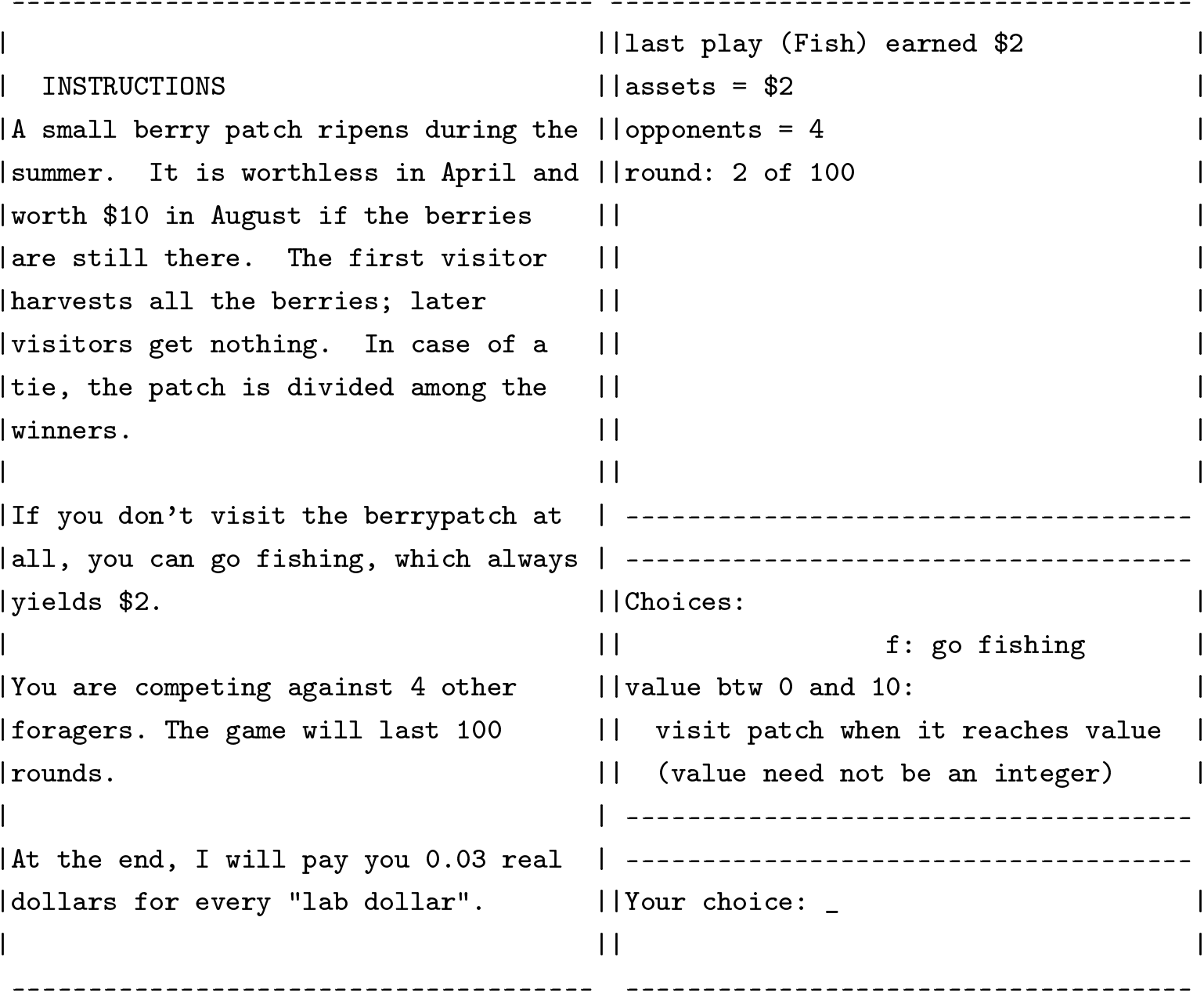
Computer screen used in berrypatch game

Subjects play in groups of five. In each round of the game, each subject chooses between “going fishing,” which yields a certain return of 2 lab dollars, and attempting to harvest the berry patch. Those who attempt the berry patch choose a value at which to harvest. A subject who chooses the value *v* will gain:

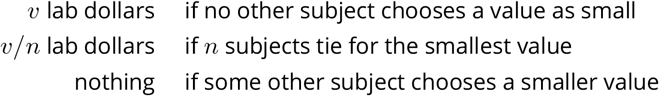

At the end of the game, subjects are paid 0.03 US dollars for each lab dollar.

Figure 2 shows the results from two experiments, each with 5 subjects, and totalling 172 trials. The students ignored the berry patch about half of the time. On those occasions when they did visit it, they were most likely to visit when the patch’s value barely exceeded the opportunity cost (the payoff from going fishing).

**Figure 2.**
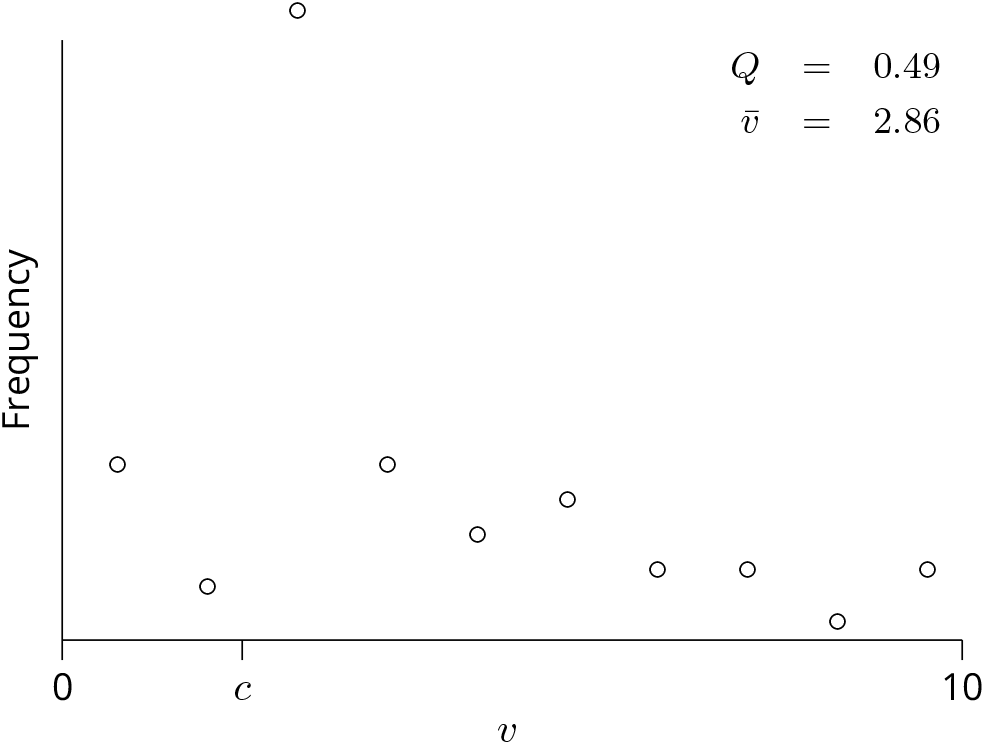
Experimental results. Data from two experiments, each with 5 subjects and totaling 172 trials. On each round of the game, subjects may choose to “go fishing” which yields reward *c* = 2. Alternatively, they may visit the berry patch when it is worth a value *v*, which they choose. Here, where 0 ≤ *v* ≤ 10. On the horizontal axis, values of *v* are grouped into 10 bins. The vertical axis shows the frequency with which the values in each bin were played. *Q* is the frequency with which “go fishing” was played, and 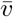 is the mean strategy among the other plays.

Now these students are not foragers, but each of them descends from a long line of foragers. It seems possible that our species has evolved a brain that is equipped to find the optimal solution to such problems. To find out whether these students reached an optimal solution, we need a model.

## 3 Model

In the experiment, the berry patch game was played 100 times. The model, however, will deal only with a single round of the game: with what is called the “stage game.” There is an easy way to justify this simplification: Because the game is played a fixed number of times, the Nash Equilibrium (NE) of the repeated game must involve repetition of that of the stage game [8, pp. 155-156]. This easy justification is suspect, however, because it depends on a feature of the game that is unrealistic. In nature animals do not compete against an unchanging group of competitors for a fixed number of rounds. In some ecological contexts, the game is played only once; in others it is played an unpredictable number of times against a variable group of competitors. Games that are repeated a fixed number of times are evolutionary novelties, and we may have evolved no adaptation to them. Nonetheless, it is always best to start simple, so this paper will deal with a single repetition of the stage game. The results will be relevant to games that are played only once, but should be applied with caution to repeated games.

In the model, *K*+1 foragers want a single resource, but that resource benefits only the foragers who first attempt to harvest it. The value *v* of the resource increases from 0 at the beginning of the season to a maximum of *V* as the resource ripens. Foragers who attempt to harvest the resource must decide how ripe to let the resource get before attempting to harvest it. The first forager to visit the resource gets it and thus gains *v*, its value when harvested. Those who try to harvest the resource later get nothing. If *n* individuals arrive at the same time, then each has an equal probability of success so that the expected payoff is *v/n*. Those who ignore the resource altogether can engage in some other activity that yields a certain payoff of *c*. I will refer to this alternative activity as “going fishing.” For notational simplicity, I set *V* = 1, which amounts to measuring all benefits and costs as proportions of V, the maximum potential benefit.

T. Bergstrom (personal communication) observes that this can also be interpreted as a model of an auction with *K* + 1 competitors. Each competitor first decides whether to pay an entry fee *c*, which allows participation in the auction. Participants then choose a bid, *b*:= *V* – *v*, and the prize goes to the highest bidder. In the literature on auctions, most authors have been concerned either with the case in which each participant values the prize differently but knows only his own valuation [9] or with the case in which each participant has a private estimate of the prize’s unknown value [10]. I take a different approach here, assuming initially that the value of the resource is known with certainty and is the same for all competitors. Computer simulations (described in the supplementary materials) indicate that the main results are not sensitive to this assumption.

These interactions are assumed to take place within some large population. Each generation, the members of the population are randomly divided into groups of size *K* + 1, and each group then plays the berry patch game. In evolutionary game theory, we are interested not in the payoff to some strategy within a particular group, but in the average payoff to that strategy across the population.

### 3.1 Pure strategies

An *evolutionary stable strategy* (ESS) is a strategy that resists invasion by all alternative strategies [7]. In the present model, no pure strategy can be an ESS. To see why, first note that it never pays to choose *v* < *c* because one can always do better than this by going fishing. Suppose therefore that nearly all of the population plays the strategy labeled *v*_0_ in figure 3. Since everyone in each group is playing the same strategy, the benefits are divided *K* + 1 ways and each individual earns *v*_0_/(*K* + 1). A rare mutant who played *v* = *v*_1_ (where *v*_1_ < *v*_0_) would always beat its neighbors to the berry patch. When rare, the mutant almost always occurs in groups by itself and therefore does not have to share the resource. Consequently, it will earn *v*_1_. It will increase in frequency when rare provided that *v*_1_ > *v*_0_/(*K* + 1). But this is always true if *v*_1_ is sufficiently close to *v*_0_. Thus, any pure strategy between *v* = *c* and *v* = 1 can be invaded by mutants playing a slightly smaller value of *v*. The strategy *v* = *c* is not an ESS either, because each member of a population playing this would earn *c*/(*K* + 1) and could do better by going fishing. The only other pure strategy is “go fishing,” which earns a payoff of *c*. But fishing is not an ESS either, for a population of fishers could be invaded by mutants playing *v* > *c*. There are thus no symmetrical equilibria in pure strategies.

**Figure 3.**
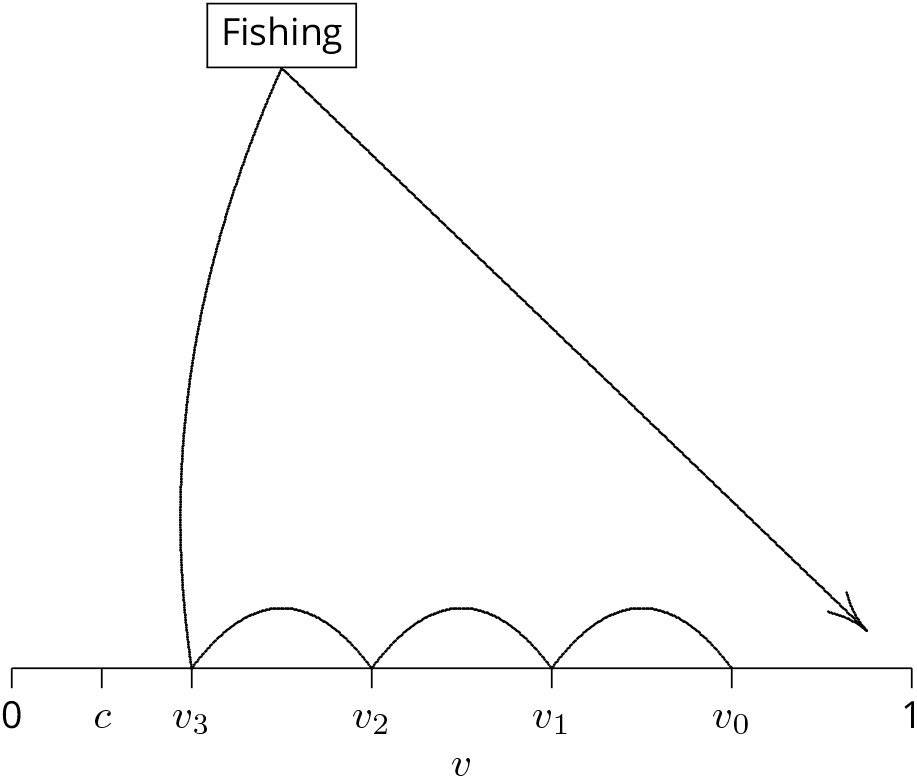
Dynamics with pure strategies

There do not seem to be any asymmetrical equilibria either. For example, suppose that within each group individual A plays *v* = *c* and the rest go fishing. Then each individual earns *c*, yet this is not a Nash equilibrium because A could do better by playing some *v* greater than *c*. Even if this were an equilibrium, it would pose a coordination problem. Asymmetrical equilibria are feasible only if there is some means of deciding in advance which player will play which role. Thus, it is of interest to consider the case in which asymmetrical equilibria are impossible, and there are no pure-strategy equilibria at all.

If there is no equilibrium in pure strategies, how can we expect the population to behave? One possibility is that the dynamics will be cyclical as suggested in figure 3. There, the population is initially fixed at *v* = *v*_0_, but the value of *v* gradually declines as successively smaller mutants invade. A mutant with strategy *v* receives payoff *v* when rare and *v*/(*K* + 1) when common. Eventually, *v* falls so low that this latter payoff is less than *c*, the payoff from going fishing. Fishing therefore increases in frequency, as indicated by the path from *v*_3_ to “fishing” on the figure. This increase continues until the resource is rarely harvested, and mutants playing large *v* can invade. This brings us back to our starting point. This story suggests that foraging behavior might exhibit cyclical dynamics, a point to which we will return.

### 3.2 A mixed equilibrium

But there is another possibility: foragers may randomize their strategies by choosing *v* from a probability distribution. Let *I* denote a strategy that chooses value *v* = *x* with probability density *f*(*x*) and chooses not to visit the resource at all with probability *Q*. I assume that all values chosen by *I* fall within an interval [*L, U*], where *c* ≤ *L* < *U* ≤ 1. In otherwords, *L* is the lowermost value ever chosen and *U* the uppermost.

The value of *L* is easy to determine. Suppose that *L* > *c*. Then a population playing *I* could be invaded by a mutant playing a fixed value of *v* that lay between *c* and *L*. Thus, *I* can be an ESS only if *L* = *c*. Analogous arguments show that *f*(*x*) > 0 over the entire interval *c* < *x* < *U* and that *f* must be a “pure density”: it cannot give non-zero probability to any point in this interval. Before determining the value of *U*, I must first derive some formulas.

The probability density *f*(*v*) is closely related to two other functions. The *survival function s*(*v*) is the probability that a forager playing strategy *I* will not have visited the resource by the time its value is *v*. It equals

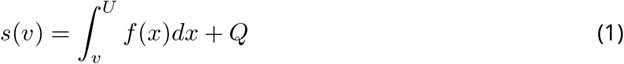

The *hazard function h*(*v*) is the conditional probability density of a visit when the resource has value *v*, given that no visit has yet been made. For convenience, I record here a series of relationships among these functions, which are well-known within demography and survival analysis [5, p. 6].

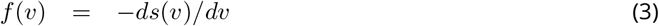

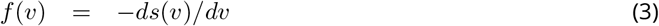

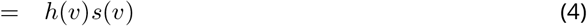

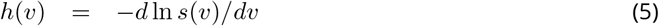

I denote by Π(*v, I^K^*) the payoff to a forager playing pure strategy *v* against *K* opponents playing *I*. This payoff equals

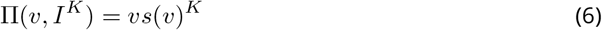

since *s*(*v*)^*K*^ is the probability that none of the *K* opponents visit the resource by the time its value is *v*. The payoff to the mixed strategy *I* is an expected value:

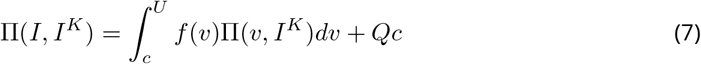

To find a formula for *s*, I make use of the fact that if *I* is an ESS, then

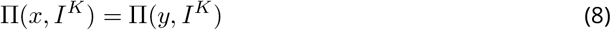

for any two pure strategies *x* and *y* that are played by I with positive probability [2, theorem 1]. In other words, all strategies receive the same payoff when playing against *I*. Consequently, the graph of Π(*v, I^K^*) against *v* must be flat, and *d*Π(*v, I^K^*)/*dv* = 0. This implies that

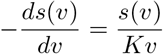

The left-hand side of this expression above equals *f*(*v*) (equation 3). Consequently, the righthand side must equal *h*(*v*)*s*(*v*) (equation 4), and the hazard function is

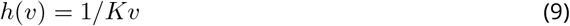

Substituting into equations 2 and 4 gives the survival and density functions:

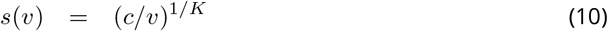

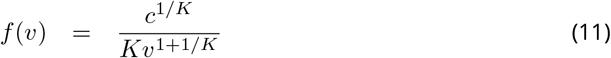

A forager will fail to visit the resource with probability

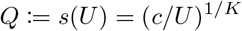

I assume that *c* > 0: that fishing pays, so that visiting the resource entails an opportunity cost. This requires that *Q* > 0—that foragers go fishing at least occasionally. Since fishing is part of the mixed equilibrium, the return from attempting to harvest the berry patch must on average equal the return *c* from fishing. On average, therefore, the net benefit from foraging must equal *c*, the opportunity cost. This insight can be verified by substituting equation 10 into equation 6, which yields Π(*v, I^K^*) = *c* irrespective of *v*. Since each pure strategy yields payoff *c* against *I*, it follows that an individual playing *I* against *I* will earn *c* too.

We are now in a position to determine the value of *U*, the uppermost pure strategy that is ever played by *I*. Consider the fate of a rare mutant playing *v* = 1 against a population playing *I*. The mutant’s fitness is

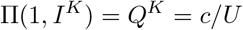

Meanwhile, the *I*-strategists each earn Π(*I, I^K^*) = *c*. The mutant’s fitness is greater unless *U* = 1. Thus, *U* must equal 1 if *I* is an ESS. Strategy *I* chooses pure strategies from the entire interval between *c* and 1; it ignores the resource with probability

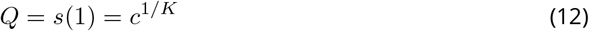

This result apparently holds in contexts more general than the present model, for it has also been derived in related models of auctions with entry fees [6, Eqn. 9]. The mean value of *v* among foragers who visit the resource is

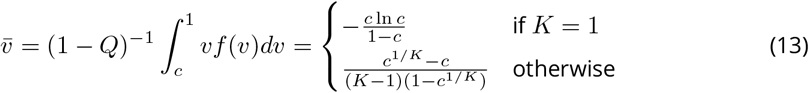

### 3.3 Stability of the mixed equilibrium

We have just seen that when everyone plays *I*, all strategies receive equal payoffs. This guarantees that *I* is a Nash equilibrium but not that it is evolutionarily stable. *I* resists invasion only if the fitness of any alternative strategy would decline as its frequency increased. To express the condition under which this is true, we need notation for payoffs against a heterogeneous mixture of opponents. Let Π(*x, J*^1^*I*^*K*−1^) denote the payoff to some strategy *x* against *K* opponents of whom 1 plays strategy *J* and *K* – 1 play *I*. Appendix section A shows that *I* resists all invasions if

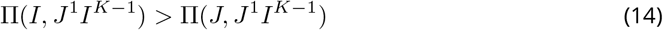

for all (pure or mixed) strategies *J* that differ from *I*. This is the multi-player analog of inequality 3 of Bishop et al. [2, p. 90]. Section A.1 shows that *I* resists invasion by all pure strategies when

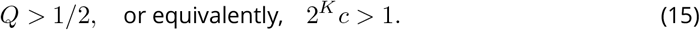

The NE resists pure-strategy invaders only if foragers are more likely to ignore the berry patch than to try to harvest it. The larger the values of *c* and *K*, the more likely this is to be so. On the other hand, the NE *never* resists invasion by mixed strategies that are similar to the NE (section A.2). Consequently, the NE is evolutionarily unstable, and this game has no ESS.

To explore the dynamics of this process, I turn now to computer simulations.

## 4 Computer simulations

### 4.1 Mixtures of pure strategies

Consider first a population in which each individual plays a pure strategy. Each simulation begins with all individuals playing *v* = 1. The first event in the life cycle is mutation, which assigns new strategies to one per cent of the population. Of these mutants, half become fishers and half are assigned a value of *v* chosen at random on the interval between 0 and 1. After mutation, fitnesses are assigned using the model above. Reproduction is haploid, with each individual producing offspring in proportion to her fitness.

Figures 4–5 show simulations with increasing values of 2*^K^c*. These should show increasing stability (ineq. 15), and indeed they do. In the less stable simulation (fig. 4), 2*^K^c* is well below unity, so *I* can be invaded. As the figure shows, 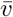 and *Q* both oscillate wildly. In figure 5, 2*^K^c* > 1, so *I* cannot be invaded by pure strategies. In this case, 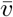 and 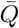 each converge rapidly toward the NE.

**Figure 4.**
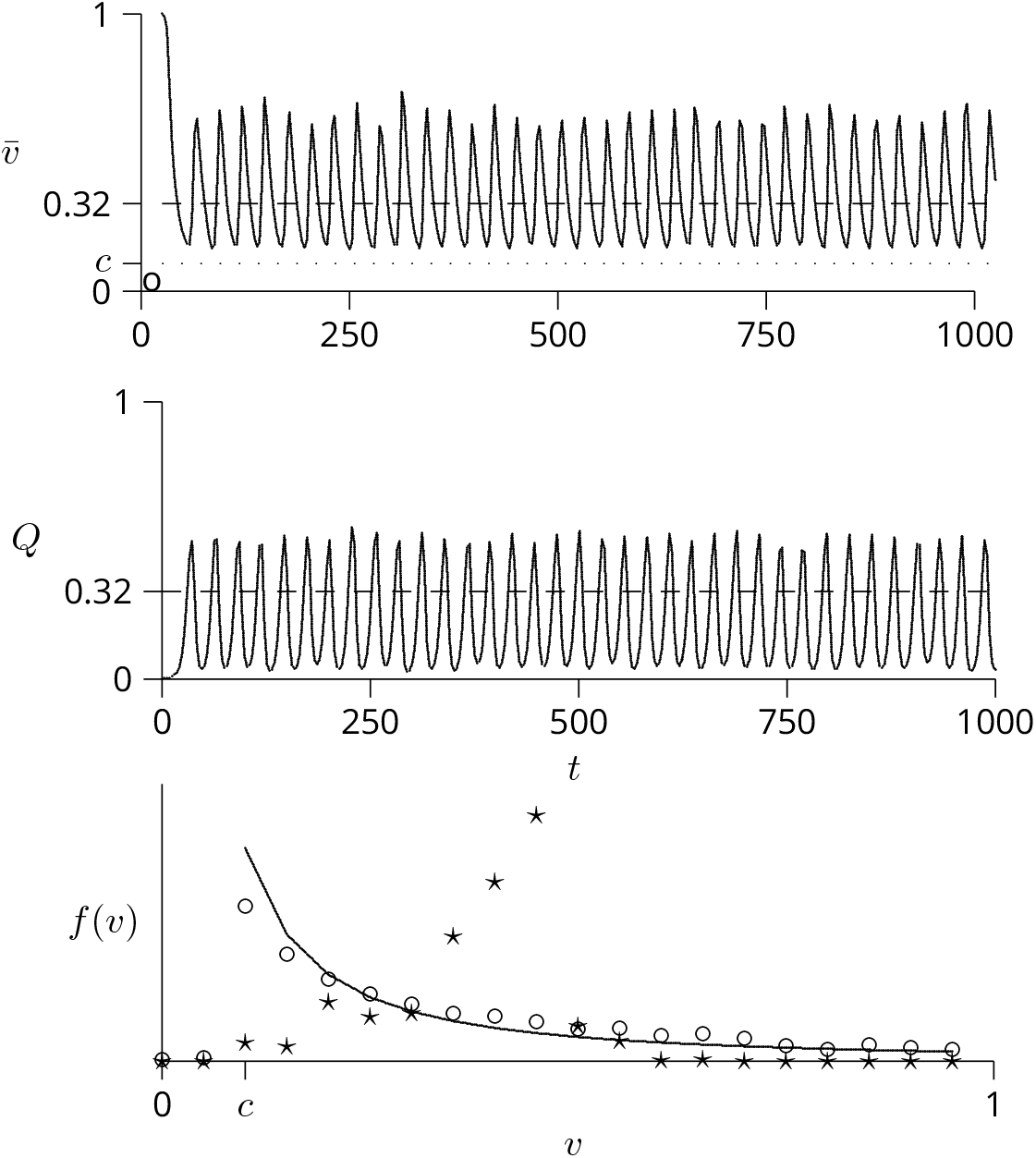
Unstable dynamics in a simulation with pure strategies, assuming *c* = 0.1, *K* = 2, and 2*^K^c* = 0.4. In this simulation 2*^K^c* < 1, so the NE does not resist invasion by pure strategies. In each generation there were 3333 groups of size 3. The dashed lines show 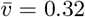 (upper panel) and *Q* = 0.32 (middle panel), the values predicted by the NE (equations 13 and 12). In the lower panel, the stars and circles show empirical frequency distributions of the strategy variable *v*. The distribution shown with stars was calculated from the simulation’s final generation, while that shown with circles aggregates over a large number of generations—all generations since the first in which v fell to the value predicted by equation 13. The solid line shows the Nash equilibrium.

**Figure 5.**
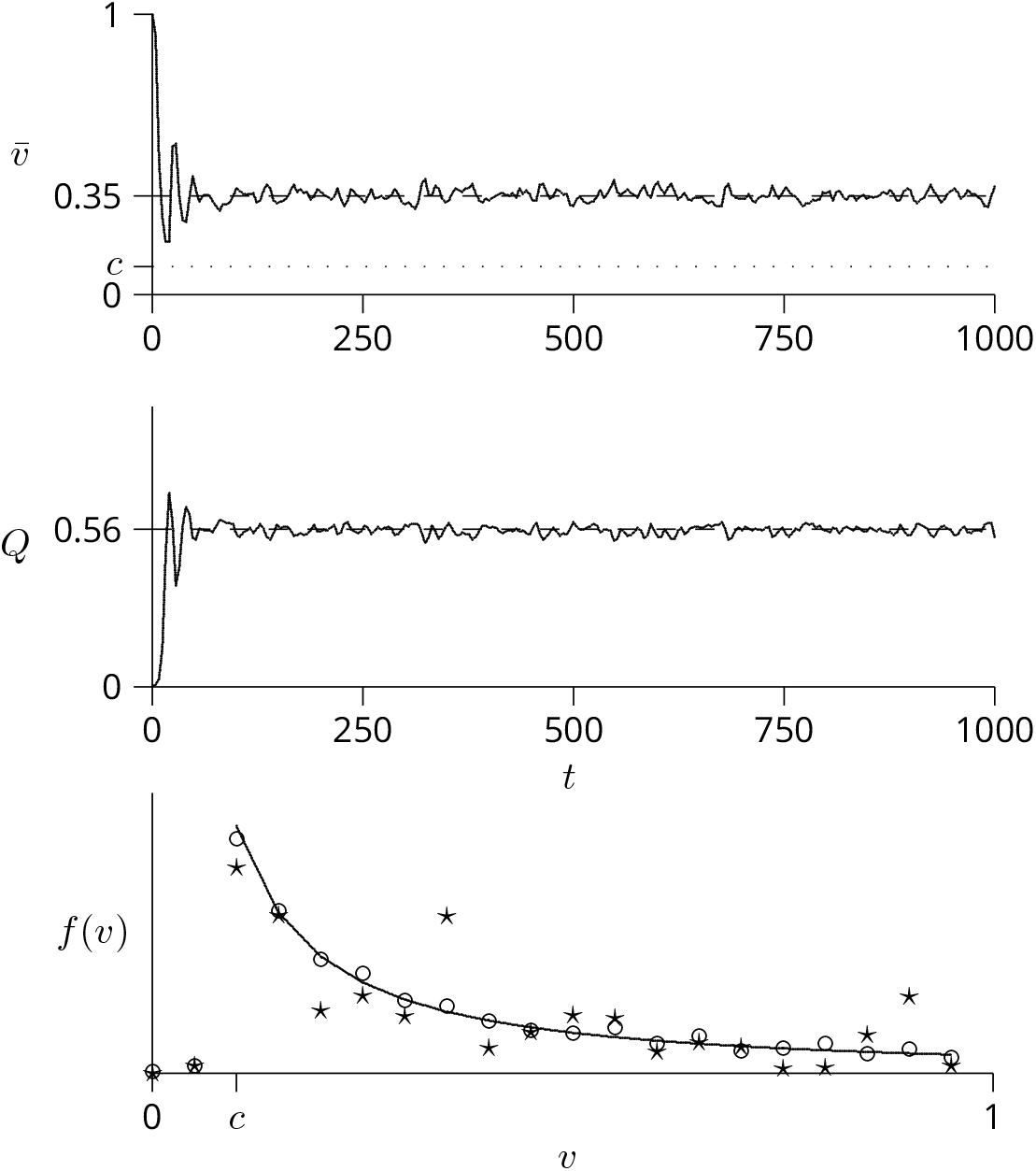
Stable dynamics in a simulation with pure strategies, assuming *c* = 0.1, *K* = 4, and 2*^K^c* = 1.6. In this simulation, 2*^K^c* > 1, so the NE resists invasion by pure strategies.

In the lower panels of these figures, the stars and circles represent empirical frequency distributions of the strategy variable *v*. The stars show distributions calculated from each simulation’s final generation, while the circles show a distribution averaged over many generations. The solid line shows the predicted frequencies at the NE, as calculated from equation 10. The starred distribution fits the NE poorly in figure 4 but fairly well in figure 5. The long term average distribution (shown by circles) fits the NE well in both cases.

Additional simulations (not shown) confirm this pattern: the system oscillates when 2*^K^c* < 1 but converges when 2*^K^c* > 1, as predicted by inequality 15.

### 4.2 Simulations with mixed strategies

Now suppose that each individual plays one of three mixed strategies, of which one is the NE and the other two are perturbed away from the Nash. To generate a perturbed strategy, I divide the interval [*c*, 1] into two segments of equal length. Within each half of this interval, the hazard is the Nash hazard (Eqn. 9) times a multiplier that is drawn independently and at random from a gamma distribution with mean 1 and variance 2.

At the beginning of the simulation, individuals are assigned the Nash strategy with probability 0.99. Otherwise, they are assigned one of the two perturbed strategies, chosen at random. In each generation, there are 2000 groups of size *K* + 1. Each individual chooses a strategy by sampling from her own mixed strategy and then plays the berry patch game with the other members of her group. The fitness of an individual equals her payoff in this game. The offspring generation is formed by sampling parents at random with replacement, weighted by parental fitnesses. The final step in each generation is mutation, which affects 1% of individuals per generation. When an individual mutates, it adopts a different strategy, chosen at random from among the other two.

Figure 6 shows the results of one simulation, in which *c* = 0.3 and *K* = 4. For these parameters, 2*^K^c* = 4.8, so condition 15 implies that the Nash equilibrium would resist invasion by pure strategies. Yet mixed strategies can clearly invade. The non-Nash strategies initially rise in frequency and then settle down to relatively stable values. Figure 7 shows the first 2500 generations of this simulation as ternary plot. After the first few generations, the strategy frequencies are constrained within a small region. Although there are no obvious cycles, it is impossible to tell whether the dynamics are cyclical or chaotic. Cycles may be obscured by the stochasticity of the simulation.

**Figure 6.**
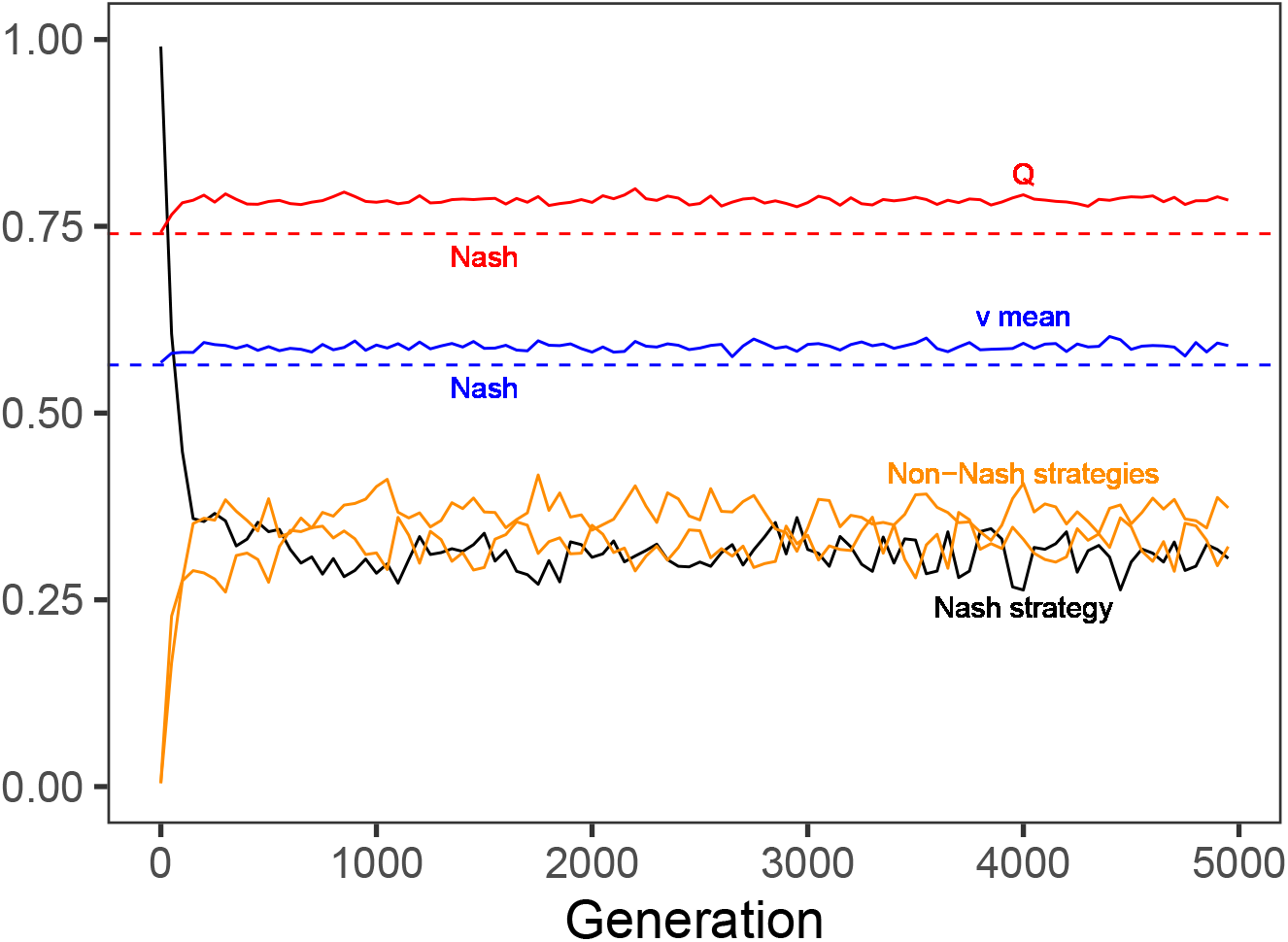
Simulation of mixed strategies, with *c* = 0.3 and *K* = 4. The black line shows the frequency of the Nash strategy, and the orange lines show those of two mixed strategies that differ from the Nash. The red line shows the frequency, *Q*, with which individuals went fishing, and the blue line shows the mean value, 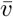, of the berry patch at the time it is harvested. The two dashed lines show the values of *Q* and 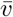 at the NE. The population consisted of 2000 groups.

**Figure 7.**
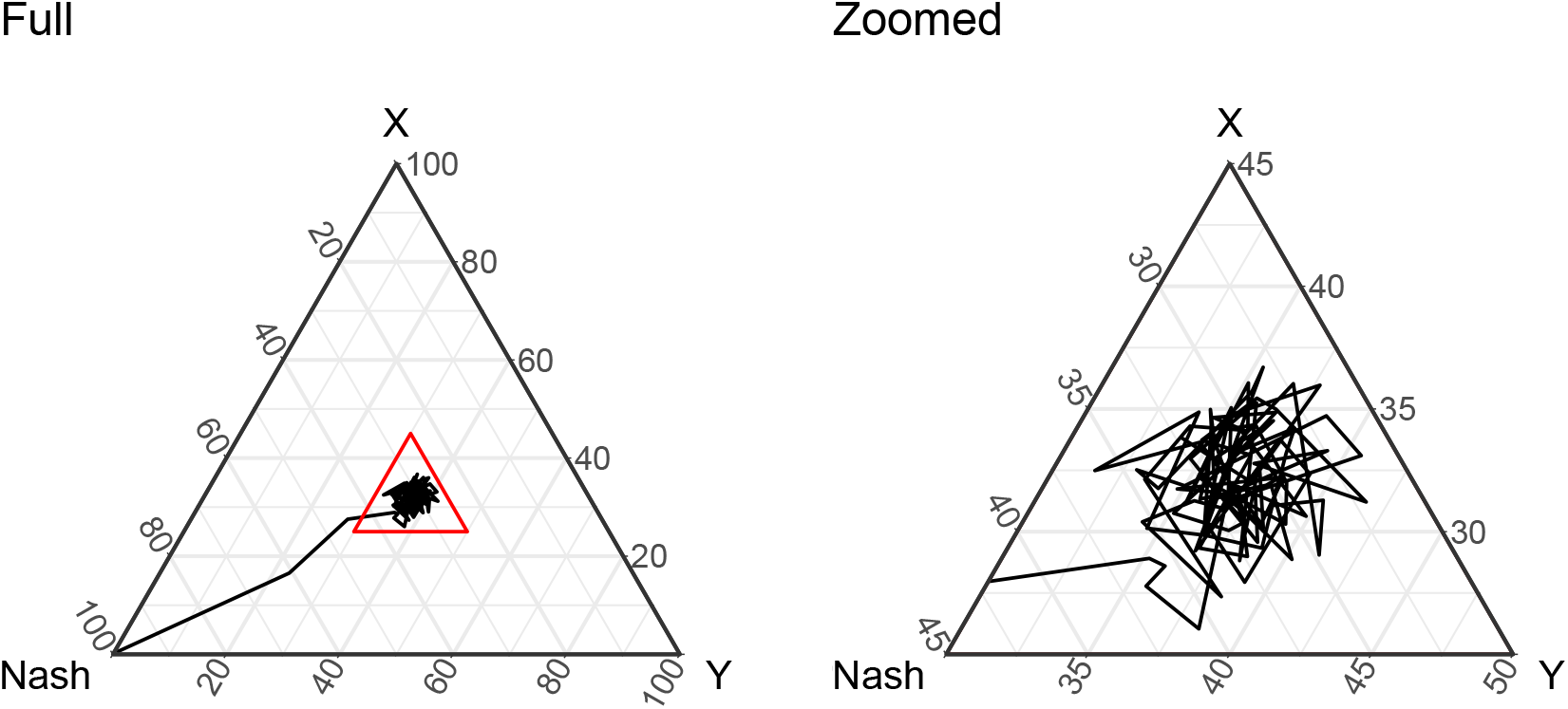
Ternary plot of the first 2500 generations of the simulation in Fig. 6. *X* and *Y* are the two non-Nash strategies. The right panel zooms in on the region of the red triangle in the left panel. Every 50th generation is plotted.

In spite of the instability of the NE, the red and blue lines in Fig. 6 are not far from the values (*Q* ≈ 0.74 and 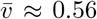) it predicts. This suggests that the NE may be predictive even though it is not an ESS. Fig. 8 supports this idea. It plots the root mean squared deviation (RMSD) from the NE against *c* and *N*. Simulations that remain near the NE have small values of the RMSD. For example, RMSD equals 0.089 for the simulation in Fig. 6. This is among the smaller values in Fig. 8, and this small value is consistent with the fact that *Q* and 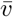 remain near the NE values throughout the simulation in Fig. 6. Figure 8 shows that the RMSD declines with *c* and also with *K*. Furthermore, the spread of this statistic also declines. This implies that—even though there is no ESS—the process tends to stay in the neighborhood of the NE when either the opportunity cost (*c*) or the number (*K*) of competitors is large.

**Figure 8.**
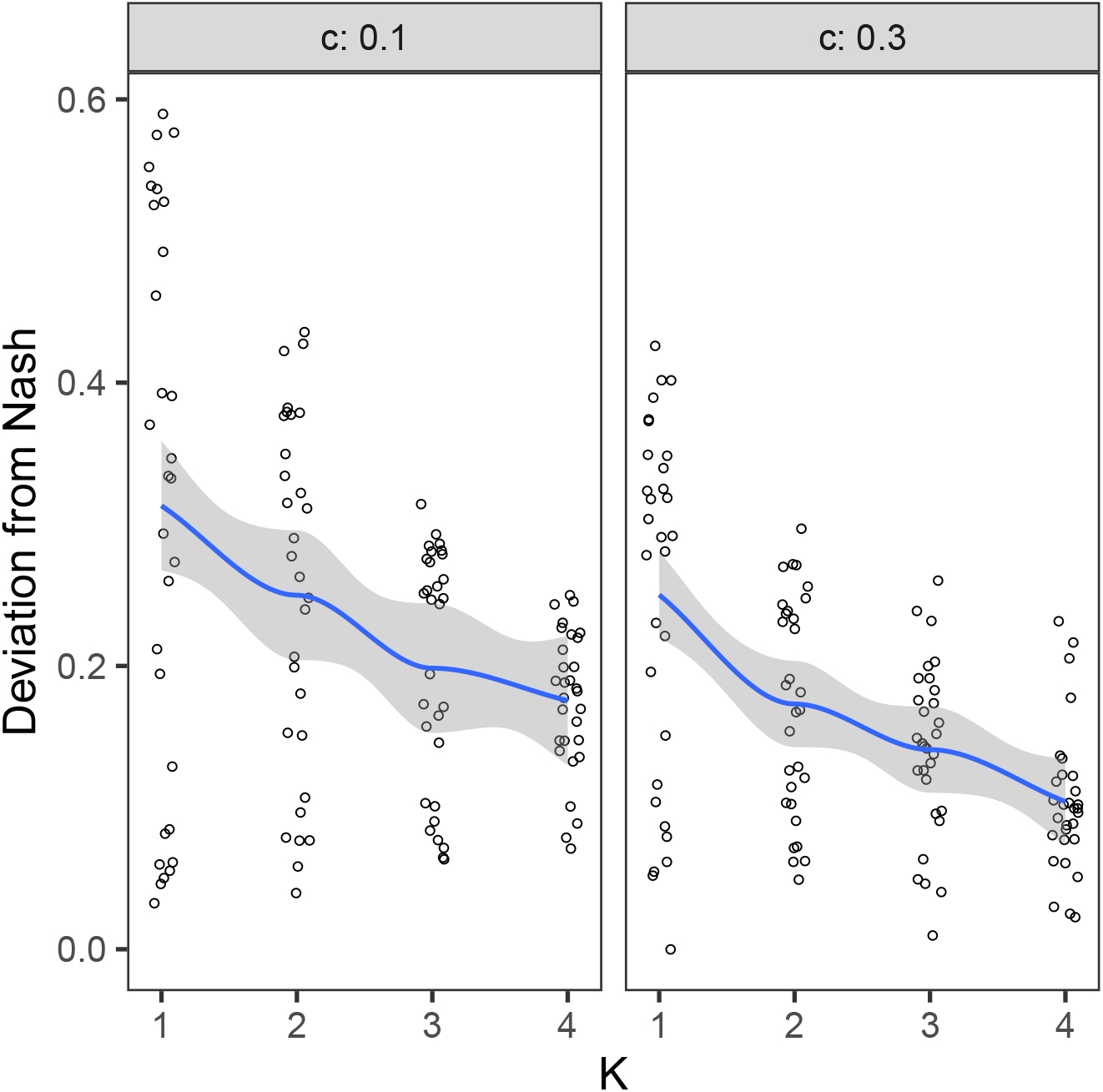
Root mean squared deviation (RMSD) from the Nash equilibrium, as a function of *K* and *c*. The mean squared deviation (MSD) of a strategy is calculated by numerically integrating the squared difference between its density function and that of the NE. The MSD of an entire simulation is the average of MSD across strategies and generations, excluding the first 1000 generations. The RMSD (plotted above) is the square root of the MSD of the simulation. Each point is a simulation of 5000 generations.

## 5 Properties of a population playing the NE

These results suggest that the NE may often be a good description of the population even though it is not an ESS. Let us therefore ask how a population that played the NE would behave. In such a population, foragers should often ignore the berry patch. This follows from the fact that the NE is predictive only when either *c* or *K* are large. In such cases, *Q* will also be large (Eqn. 12), so foragers will often go fishing rather than visiting the berry patch.

Surprising results emerge when one asks such questions as “How does the value of the harvested resource change with the number of competitors?” Intuition suggests that when the number of competitors is large, the resource will usually be harvested sooner and ata lower value. As we shall see, however, the model implies precisely the opposite.

Consider the probability distribution of the resource’s value at the time it is harvested. If *K* + 1 foragers are all playing strategy *I*, then the survival function of this new random variable is

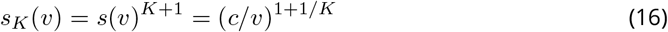

This survival function gives the probability that the resource survives unharvested at least until its value is *v*. Equations 5 and 4 now give the hazard function and probability density:

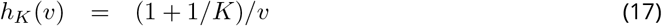

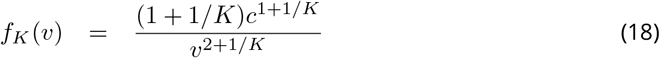

The probability that the resource is never harvested equals

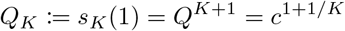

For example, if *c* = 1/2 then the resource remains unharvested 1/4 of the time with two competitors but 1/2 the time with an infinite number. Apparently (but contrary to intuition), larger numbers of foragers leave more fruit on the tree.

The probability density of the value of resource when it is harvested, given that it is harvested at all, is *f_K_*(*v*)/(1 – *Q_K_*). The mean value of the resource when harvested is therefore

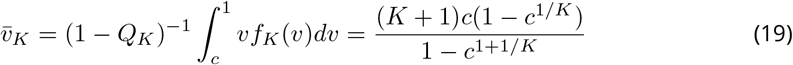

In the special cases of two competitors and of an infinite number, 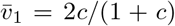 and 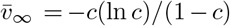. With *c* = 0.6, these two cases give 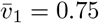 and 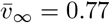. Note that 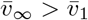. This means that on average (and contrary to intuition), the mean value of the resource when harvested increases with the number of competitors.

## 6 Return to experimental data

The experimental data are re-plotted in Fig. 9 along with corresponding theoretical results. The fit between observation and the NE is far from perfect: The observed value of *Q* is low (1/2 rather than 2/3), and the data give too much weight to values of *v* between 2 and 3. It is tempting to interpret these discrepancies in ecological terms or in terms of the differences between model and experiment. For example, the predominance of low values of *v* might result from risk aversion. Or the discrepancies might reflect the fact that the model describes a one-shot game, whereas the subjects played multiple rounds.

**Figure 9.**
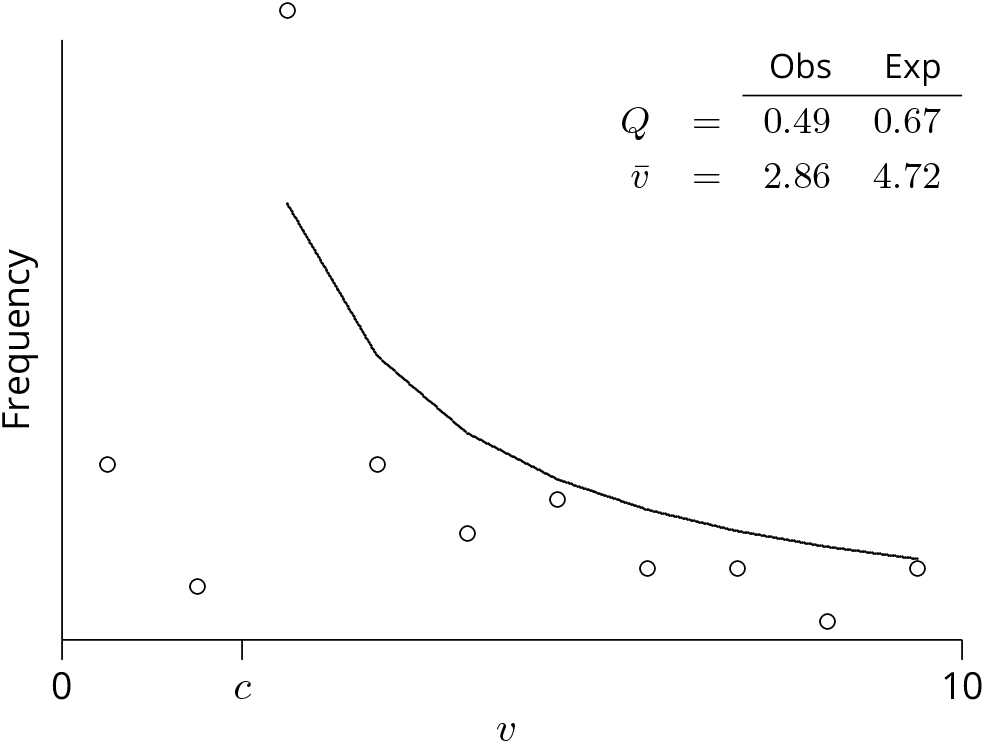
The Nash equilibrium (solid line and values under”Exp”) compared with data from Fig. 2. In the experiment, *K* = 4, *v* ranged from 0 to 10, and “going fishing” yielded payoff *c* = 2. In the model, payoffs are re-expressed as fractions of the maximum payoff, so that *c* and *v* both lie between 0 and 1.

I am reluctant, however, to interpret the results in this way. Even in the best of circumstances, we only expect populations to be near the NE, not at it. We should not expect precise numerical agreement, even when all the model’s assumptions hold. Emphasis should instead be on the model’s qualititative predictions.

In qualitative terms, there is good agreement between the NE and the results in Fig. 9. As predicted, these human subjects often ignored the berry patch. And when they did attempt to harvest, they tended to do so when the reward barely offset the cost. Although these similarities do not show that human behavior was shaped by the evolutionary process described here, they are broadly consistent with that view.

## 7 Discussion

In real berry patches, the berries do not all ripen on the same schedule, and multiple foragers make multiple trips to compete for the berries that are currently (somewhat) ripe. The present model is thus an abstraction, intended to capture the essential features of competition for a resource that increases gradually in value. It is possible that the model’s artificial features exaggerate its instability. Nonetheless, it probably captures some of what goes on in nature.

In the current model, no pure strategy can be evolutionarily stable, so any equilibrium must be mixed. Yet the only mixed Nash equilibrium is evolutionarily unstable—it does not resist invasion by other mixed strategies. Nonetheless, the dynamics of this process remain in the neighborhood of the NE if either the cost of harvesting or the number of competitors is large. In such cases, the NE provides a useful description in spite of its instability.

It does not, however, provide a precise numerical description. Because the NE is evolutionarily unstable, populations should seldom be at it, although they may often be near it. We should therefore emphasize the model’s qualitative implications rather than its numerical ones. These qualitative implications are surprising. When a population is near the NE, the resource should often go unharvested, most harvesting attempts should occur when resource is barely ripe enough to offset costs, and harvesting attempts should decline in frequency as the resource ripens. The more foragers there are, the greater the chance that the resource will go unharvested and the higher its mean value when harvested.

These conclusions apply not only to foraging but also more generally whenever there is competition for something that gradually increases in value. Paul Smaldino (personal communication) compares the berry patch to scientific publishing: the longer you work on a piece of research, the better it gets, but also the greater the chance that someone else will publish your result first.

## Acknowledgements

This work began during a sabbatical leave at the Research Centre of King’s College, Cambridge, U.K. I am grateful for comments from S. Beckerman, T. Bergstrom, R. Boyd, E. Cashdan, E. Charnov, H. Gintis, A. Grafen, R. Hames, H. Harpending, K. Hawkes, J. Hirshleifer, J.H. Jones, S. Josephson, J. Kagel, Aaron McDonald, R. McElreath, Graeme Mitchison, M. Nadler, J. O’Connell, A. Roth, Paul Smaldino, and Pontus Strimling. I am also grateful to François Munoz, François Massol, Jeremy Van Cleve, and one anonymous reviewer, who reviewed the manuscript for *PCI Mathematical and Computational Biology*. Finally, I thank those who helped by playing the berry patch game. Source code and data are available at https://osf.io/2mbq4. Version 8 of this preprint has been peer-reviewed and recommended by *Peer Community In Ecology* (https://doi.org/10.24072/pci.ecology.100088)”

## A When does *I* resist invasion?

Each forager competes in a group with *K* others. Let *P*_0_ denote the probability that a random forager competes with *K* non-mutants playing *I* and let *P*_1_ denote the probability that she competes against one mutant playing *J* and *K* – 1 non-mutants playing *I*.^1^ If *J* is rare, we can neglect the possibility that more than one opponent plays *J*; hence *P*_0_ + *P*_1_ = 1. The fitnesses of *J*-strategists and *I*-strategists are

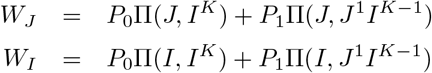

where the notation Π(*s, J*^1^*I*^*K*−1^) refers to the expected payoff to strategy *s* when playing against *K* competitors, of whom 1 is playing *J* and the other *K* – 1 are playing *I*. Strategy *I* is an ESS if and only if *W_I_* > *W_J_*. The definition of *I* implies that Π(*J, I^K^*) = Π(*I, I^K^*) [2, theorem 1]. It follows that *W_I_* > *W_J_* if and only if

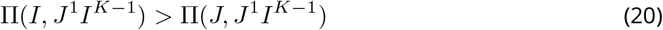

### A.1 Invasion by pure strategies

This section shows that the NE resists invasion by all pure-strategy invaders if and only if *Q* > 1/2. I assume that pure strategists either fish or play *v* ∈ [*c*, 1], because strategies *v* < *c* are dominated by the fishing strategy.

Consider first the payoff to a mutant that plays pure strategy *v* in a group with one other mutant. The two mutants playing pure strategy *v* beat the *K* – 1 *I*-strategists to the resource with probability *s*(*v*)^*K*−1^, in which case they split the prize and each receive *v*/2. Thus,

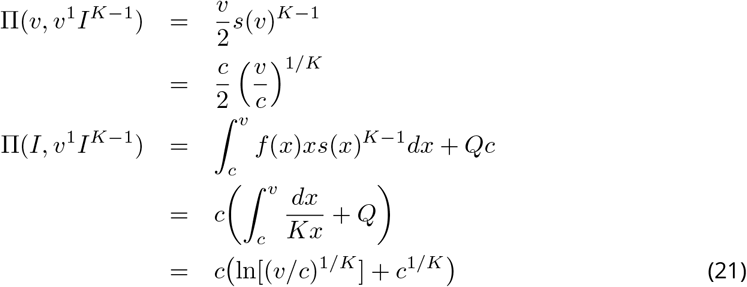

Stability requires that Π(*I, v*^1^*I*^*K*−1^) > Π(*v, v*^1^*I*^*K*−1^), which is equivalent to or

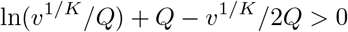

or

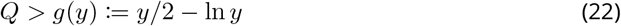

where *y*:= (*v/c*)^1/*K*^. *I* resists invasion by all *v* ∈ [*c*, 1] only if (22) holds for all *y* ∈ [1,1/*Q*]. The function *g*(*y*) has a global minimum at *y* = 2 and decreases with *y* when *y* < 2.^2^

If (22) holds for all *y* ∈ [1,1/*Q*], then it must hold for *y* = 1, in which case (22) becomes *Q* > 1/2. This condition is not only necessary but also sufficient: if *Q* > 1/2, then *y* ∈ [1,1/*Q*] ⟹ *y* < 2. This implies that *g*(*y*) decreases throughout the range of *y* and reaches its maximum at *y* = 1. When (22) holds for this maximal value, it holds everywhere. Thus, inequality 22 holds if and only if *Q* > 1/2.

It remains to consider the case of a mutant that always fishes, never visiting the berry patch at all. Let us call this strategy *F*. The payoff to *F* is always *c*, no matter what its opponents do. Strategy I will resist invasion by *F* provided that

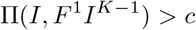

This payoff can be found by setting *v* = 1 in Eqn. 21. The resulting expression is greater than *c* for all permissible values of *c* and *K*. Thus, strategy *I* always resists invasion by *F*. This completes the justification of inequality 15.

### A.2 Invasion by a mixed strategy

This section will show that *I* never resists invasion by nearby mixed strategies. Subscripts will be used to distinguish quantities referring to different strategies: the survival and density functions of strategy *I* are denoted by *s_I_* and *f_I_*, and the corresponding functions of strategy *J* are *s_J_* and *f_J_*. The payoffs to *I* and *J* against groups with one *J* are

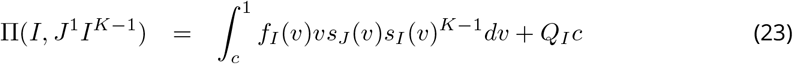

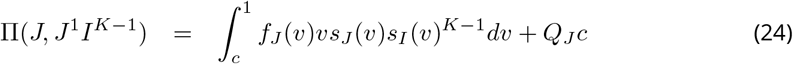

Substituting *vs_I_*(*v*)^*K*−1^ = *c/s_I_*(*v*) and 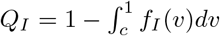 leads to

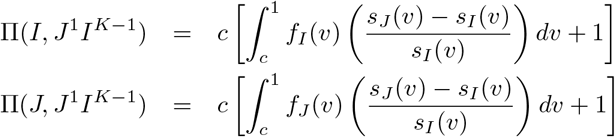

To measure the difference between the two payoffs, define

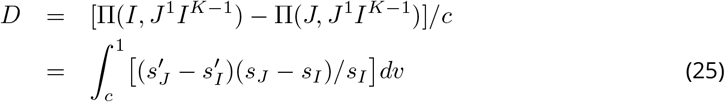

where the arguments of *s_J_* and *f_J_* have been suppressed and *f_I_* and *f_J_* have been re-expressed as 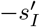 and 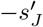 (see equation 3). To prove that *I* is evolutionarily unstable, I must show that *D* < 0 for some *J*. Now *D* = 0 when *J* = *I*, for then *D* is then the difference between two identical quantities. Consequently, I can prove that *I* is evolutionarily unstable by showing that *D* is greater when *J* = *I* than otherwise. To this end, I use the calculus of variations to show that *D* reaches a local maximum where *s_J_* = *s_I_*.

The integrand within the definition of *D* can be written as

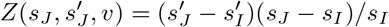

The function *s_J_* that maximizes *D* must satisfy the Euler equation [4, p. 7],

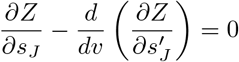

Solving this differential equation with initial condition *s_J_*(*c*) = 1 gives *s_J_*(*v*) = (*c/v*)^1/*K*^, which is identical to *s_I_*(*v*) (see equation 10). Thus, the only strategy that can possibly maximize *D* is strategy *I*, as defined in equations 9–11.

The calculus of variations requires functions with exogeneously determined endpoints. Consequently, I will stipulate that *s_J_*(1) and *s_I_*(1) are both equal to *Q_I_*, as given in equation 12. If I can show that *I* cannot resist invasion by strategies that are constrained in this fashion, then it certainly cannot resist invasion by strategies chosen without constraint.

The Euler equation provides only a necessary condition and does not guaranteee that *s_I_* maximizes *D* rather than minimizing it. To ensure that *s_I_* is indeed a minimum, one must show that the “second variation” of *D* is positive. The second variation is [4, p. 35]

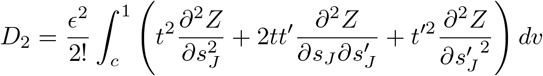

where *ϵ* is a small perturbation and *t* a function of *v* that is arbitrary except for the requirement that *t*(*c*) = *t*(1) = 0. In the present case, 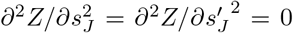, and 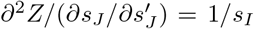. Thus, the integral in *D*_2_ becomes

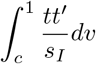

Integrating by parts produces

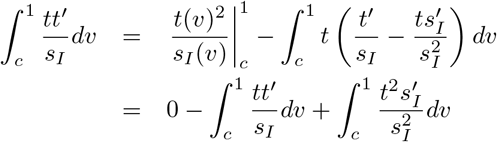

or

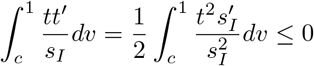

The sign of the final term follows from the observations that (*t/s_I_*)^2^ ≥ 0 and 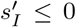 because survival functions cannot increase. This indicates that the second variation is negative, which implies that *s_I_* maximizes *D*. This shows that *I* is not an ESS. It can be invaded by any mixed strategy that is sufficiently similar to it.

## 1 Simulations with uncertainty

The theory presented in the main text [3] assumes that all foragers value the resource equally, and none have any private information about its value. To investigate the model’s sensitivity to these assumptions, I compared it against two simulations.

Each simulation mimics a model drawn from the literature on auctions [2]. In the *common value* model, the value of the resource is not known with certainty, but each forager has an independent estimate. For example, when hummingbirds compete for nectar from a flower, each bird knows the time of its own last visit to the flower but does not know about visits by other birds [1]. Thus, each bird has a private signal about the flower’s quality, but no bird knows the quality with certainty. In the *private values* model, the caloric value of the resource is common knowledge, but each forager values these calories differently. For example, a starving forager and one well fed might value the same strawberry very differently.

To model these interactions, I assume as before that foragers choose a value *v* at which to visit the resource and that the forager with the lowest value gets the resource. The return to a successful forager, however, is not *v*, but *Yv*, where *Y* is a random variable with a distribution that is uniform on the interval [0.7,1.3). This introduces a substantial uncertainty, since the range of *Y* is 60% of its mean. In these models, *v* is no longer the value of the resource at the time of the forager’s visit. I will refer to it as the resource’s “*ex ante* value” to indicate that it is the expected value before any private signal is received. In the private values model, an independent value of *Y* is drawn for each forager, and this value is revealed to the forager as a private signal. In the common value model, an independent value of *Y* is drawn for each group of *K* + 1 competing foragers, and each forager receives a signal *s* = *uY*, where *u* is uniform on [0,1) and an independent value of *u* is drawn for each forager. In either model, each forager determines its behavior by consulting its own “response function.”

Response functions are piece-wise linear with six parameters. Figure S1 shows the response functions of two foragers, one indicated by open circles and the other by bold dots. The signal is on the horizontal axis. The vertical positions of the symbols indicate the values of the six parameters in each forager’s response function. Each function is constructed by connecting the symbols with straight lines. When the value of the response function exceeds 1.3 (the maximum value that the resource can have), the forager does not visit the resource. Thus, the forager whose function is indicated by open circles never visits the resource at all. When the response is less than 1.3, its value is taken to equal *v*, the *ex ante* value of the resource at the time of the forager’s visit. Thus, the forager indicated by bold dots always visits the resource. Mutation acts by perturbing the value of each parameter in the response function. Offspring inherit the response functions of their (single) parent.

**Figure S1:**
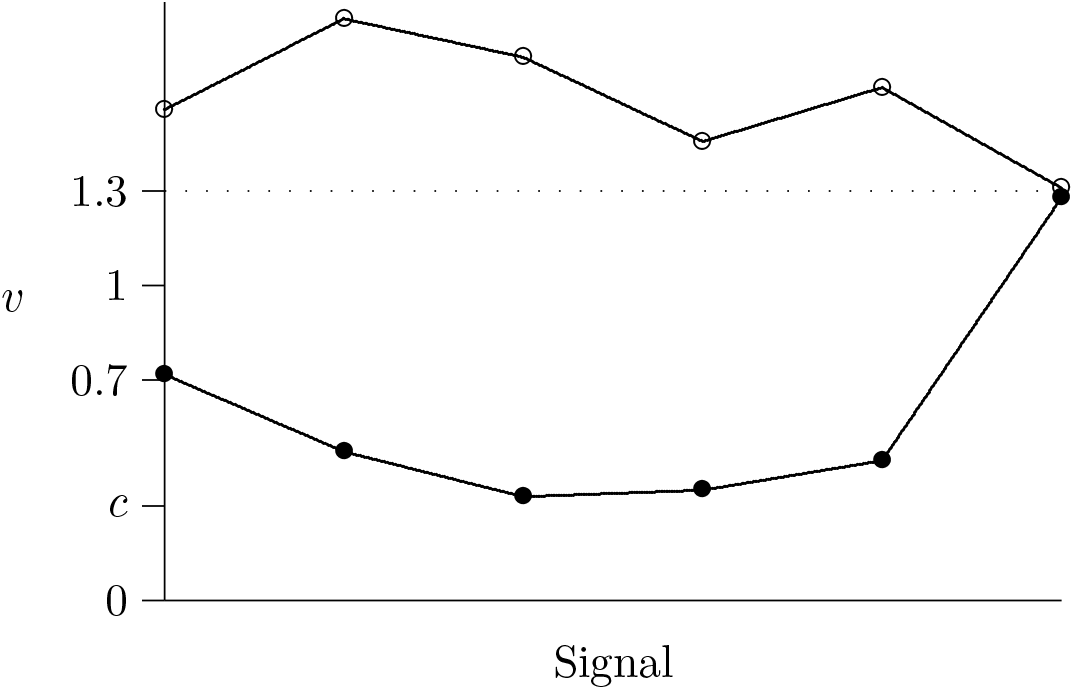
Piecewise linear response functions. The private signal is on the horizontal axis, and the forager’s response *v* is on the vertical axis. When *v* ≥ 1.3, the forager does not visit the resource and therefore pays no cost. When *v* < 1.3, the forager visits the resource when its *ex ante* expected value is v. The function indicated by open circles describes a forager who never visits the resource, and the function indicated by bold dots describes a forager who always visits the resource.

Figures S2 and S3 show the results of simulations under the common value model and the private values model, respectively. In each case, the parameter values are identical to those in figure 5 Stable dynamics in a simulation with pure strategies, assuming *c* = 0.1, *K* = 4, and 2*^K^c* = 1.6. In this simulation, 2*^K^c* > 1, so the NE resists invasion by pure strategies.figure.caption.5 of the main text. The theory presented in the main text ignores the uncertainty inherent in these simulations but nonetheless does a good job of predicting their behavior. The fit of the common value model could hardly be better. Under the private values model, the aggregate distribution fits much better than the distribution of the final generation. Nonetheless, the theoretical model is remarkably accurate even where its assumptions are not met.

**Figure S2:**
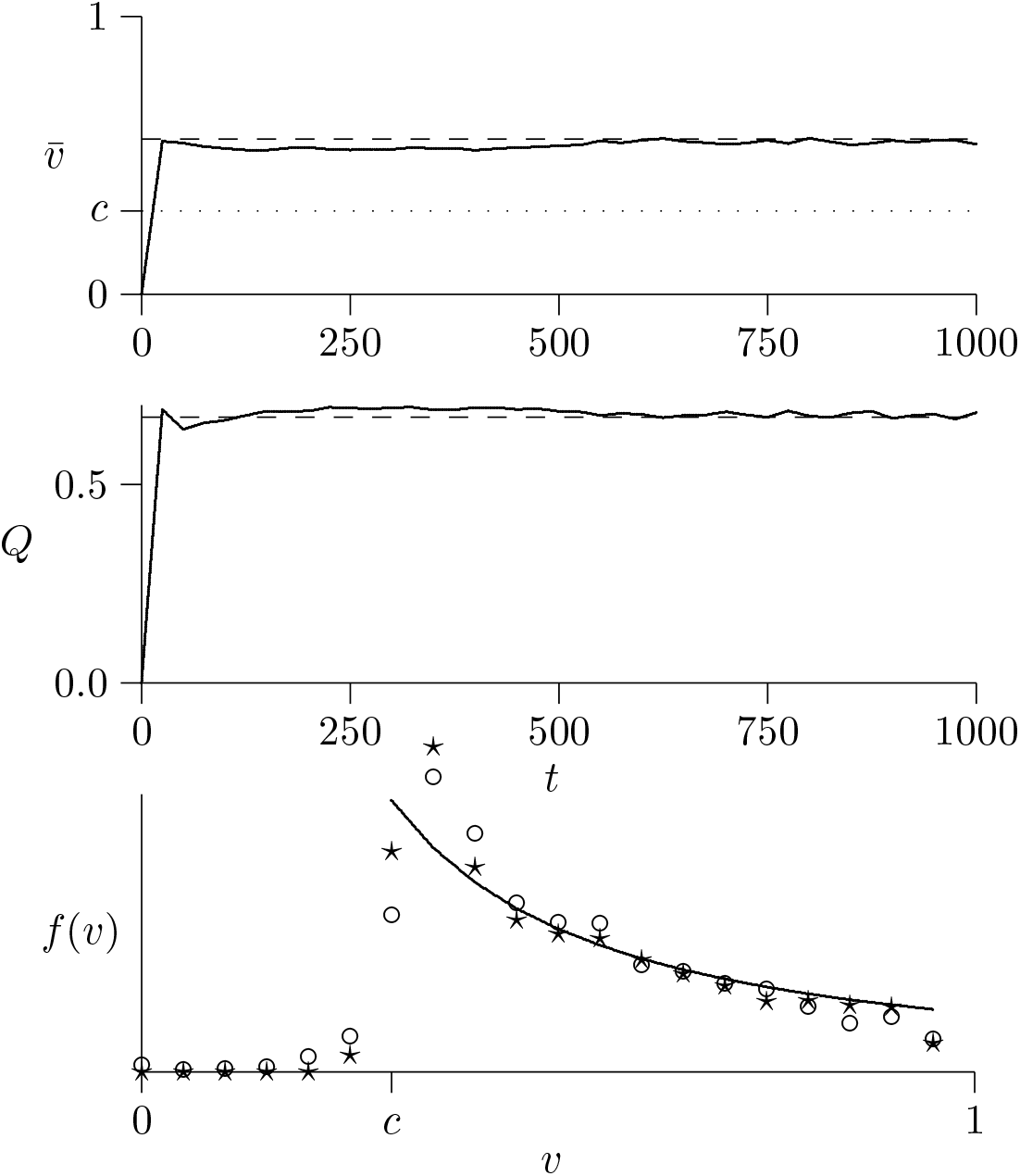
Simulation under the common value model. The simulation uses 5000 groups and assumes *c* = 0.3 and *K* = 3, so 2*^K^c* = 2.4 > 1, and the Nash equilibrium resists invasion by pure strategies. In the bottom panel, the aggregate distribution (circles) summarizes 6,530,751 individuals, and the final distribution (stars) summarizes 6386.

**Figure S3:**
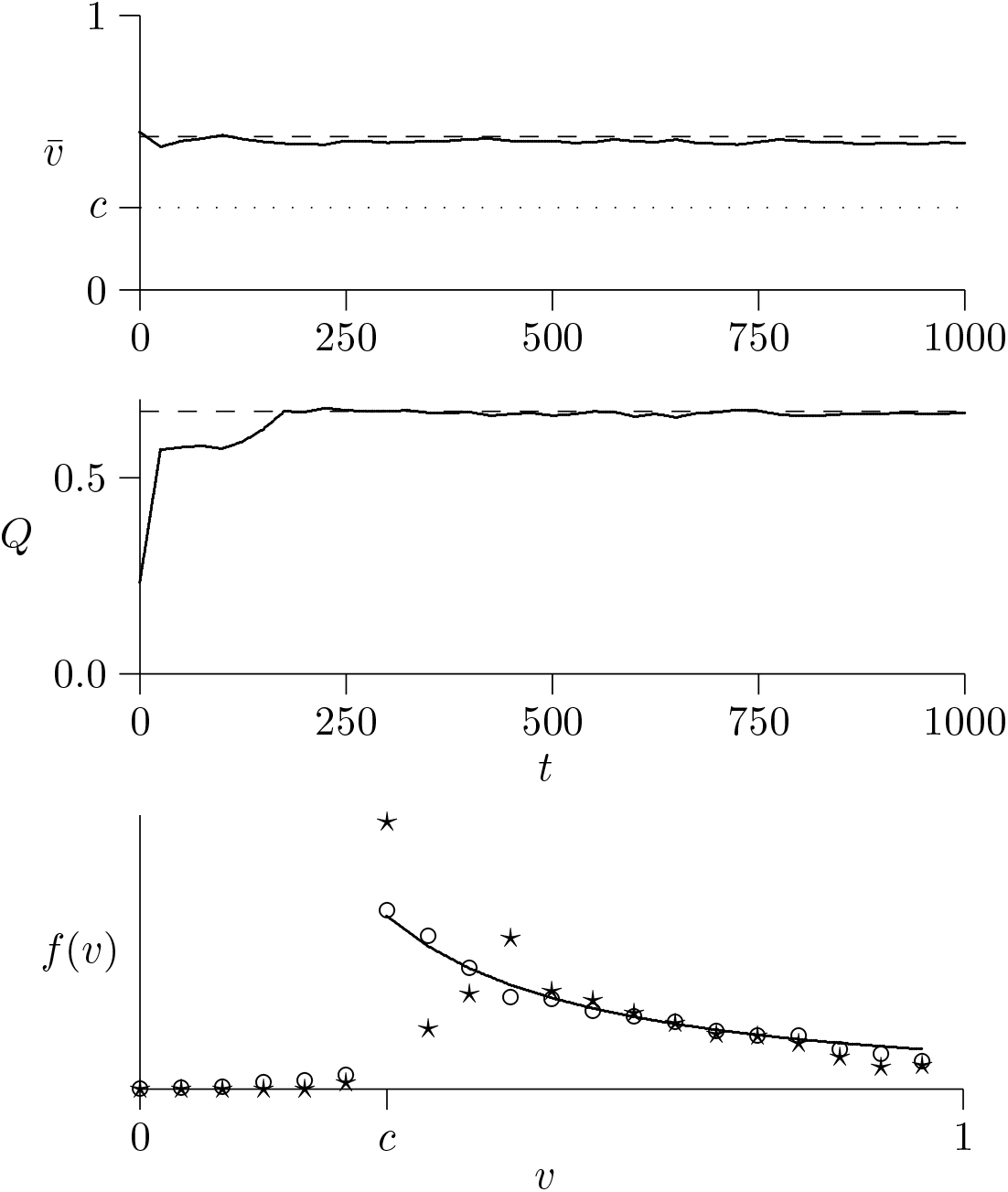
Simulation under the private value model. The simulation uses 5000 groups and assumes that *c* = 0.3 and *K* = 3, so the Nash equilibrium resists invasion by pure strategies. In the bottom panel, the aggregate distribution (circles) summarizes 6,759,231 individuals, and the final distribution (stars) summarizes 6695.

## Footnotes

1 In a finite population, *P*_0_ and *P*_1_ would be different for a *J* strategist than for an *I* strategist. These differences are small however in large populations, and they disappear in infinite ones.

2 The first derivative, *g′*(*y*) = 1/2 – 1/*y*, equals zero only at *y* = 2. The second derivative, *g″*(*y*) = *y*^−2^ > 0, is positive.

